# Mapping QTL for spike fertility related traits in two double haploid wheat (*Triticum aestivum L.*) populations

**DOI:** 10.1101/2020.10.08.331264

**Authors:** Nicole Pretini, Leonardo S. Vanzetti, Ignacio I. Terrile, Guillermo Donaire, Fernanda G. González

**Affiliations:** Centro de Investigaciones y Transferencia del Noroeste de la Provincia de Buenos Aires (CITNOBA, CONICET-UNNOBA-UNSADA). Monteagudo 2772 CP 2700, Pergamino, Buenos Aires, Argentina; Instituto Nacional de Tecnología Agropecuaria (INTA). EEA INTA Marcos Juárez. Ruta 12 s/n CP 2850, Marcos Juárez, Córdoba, Argentina; Consejo Nacional de Investigaciones Científicas y Técnicas (CONICET). Godoy Cruz 2290 CP C1425FQB, Buenos Aires, Argentina; Instituto Nacional de Tecnología Agropecuaria (INTA). EEA INTA Pergamino. Ruta 32, km 4,5 CP 2700, Pergamino, Buenos Aires, Argentina

**Keywords:** grain number, grain weight, spike length, spikelets per spike, fertile florets, chaff

## Abstract

In breeding programs, the selection of cultivars with the highest yield potential consisted in the selection of the yield *per se*, which resulted in cultivars with a higher grain number per spike (GN) and occasionally higher grain weight (GW) (main numerical components of the yield). This task could be facilitated with the use of molecular markers such us single nucleotide polymorphism (SNP). In this study, quantitative trait loci (QTL) for GW, GN and spike fertility traits related to GN determination were mapped using two double haploid (DH) populations (Baguette Premium 11 x BioINTA 2002 and Baguette 19 x BioINTA 2002, BP11xB2002 and B19xB2002). Both populations were genotyped with the iSelect 90K SNP array and evaluated in four (BP11xB19) or five (B19xB2002) environments. We identify a total of 305 QTL for 14 traits, however 28 QTL for 12 traits were considered significant with an R^2^ > 10% and stable for being present at least in three environments. There were detected eight hotspot regions on chromosomes 1A, 2B, 3A, 5A, 5B, 7A and 7B were at least two major QTL sheared confident intervals. QTL on two of these regions have previously been described, but the other six regions were never observed, suggesting that these regions would be novel. The R5A1 (*QSL.perg-5A, QCN.perg-5A,QGN.perg-5A)* and R5A.2 (*QFFTS.perg-5A, QGW.perg-5A)* regions together with the *QGW.perg-*6B resulted in a final higher yield suggesting them to have high relevance as candidates to be used in MAS to improve yield.

**Author contribution statement:** 

**Key message:** 28 stable and major QTL for 12 traits associated to spike fertility, GN and GW were detected. Two regions on 5A Ch., and *QGW.perg-*6B showed direct pleiotropic effects on yield.

## 1. Introduction

Wheat (*Triticum aestivum* L.) is one of the most cultivated and consuming worldwide cereals. Its production has to increase to supply the growing world population demand (FAO 2017; Borlaug 2007; Chand et al. 2009). Improving the cultivar’s yield potential (i.e., the yield of a cultivar adapted to the environment, which is growing without water or nutrient deficits and with no biotic stress, Evans et al. 1993) by breeding is a sustainable alternative to guarantee increases in world production (Fischer and Edmeades 2010; Fischer et al. 2014). Wheat breeding of yield potential has been based on empirical selection of yield *per se* due to the complexity of the character and the lack of knowledge and useful tools with real applicability in breeding programs (Snape and Moore 2007). This selection generally resulted in more grains per spike (GN), and hence, increased grains per unit area (no consistent trend in spikes per unit area were reported) (Waddington et al. 1986; Perry and D’Antouno 1989; Siddique et al. 1989; Slafer and Andrade 1989, 1993; Acreche et al. 2008; Del Pozo et al. 2014; Lo Valvo et al. 2018). The grain weight (GW) showed no change with breeding, except for some recent reports were yield potential was positively associated with its increment (Sadras and Lawson 2011; Aisawi et al. 2015; Yao et al. 2019).

The selection process could be more efficient using molecular markers. The implementation of single nucleotide polymorphism (SNP) markers in plan breeding had increased the pace and precision of plant genetic analysis, which in turn facilitated the implementation of crop molecular improvement (Mammadov et al. 2012). SNP markers have been increasingly used for QTL mapping studies because they are the most frequent variations in the genome, and they provide a high map resolution (Bhattramakk et al. 2002; Jones et al. 2007; Mammadov et al. 2012;). Therefore, the identification, understanding and incorporation of related yield QTL could be useful as selection tools for breeding programs.

The most common approaches looking for genetic bases to further improving yield potential are based on the numerical component analyses, GN and GW (see references quoted in Table S1). The GN is understand as the result of the total spikelets per spike (TS) and the grains per spikelet, being the former associated with the spike length (SL) and compactness of the spike (CN). Several QTL had been reported for the GW and the GN itself and their numerical components during last years. Many studies identified stables QTL for these traits widespread in the genome (see Table S1). However, considering the IWGSC Ref. Seq. v1.0 wheat genome assembly (Appels et al. 2018) we identified some QTL reported for the same trait that are located at the same position (Table 1). For example, a QTL for GW was detected in 6 studies in the chromosome 7A (Li et al. 2015; Wang et al. 2017; Daba et al. 2018; Li et al. 2018; Ma et al. 2018; Guan et al. 2018) (Table 1). Another two important QTL for SL were detected in chromosomes 2D and 5A (Börner et al. 2002; Wu et al. 2012; Xu et al. 2014; Zhai et al. 2016; Chen et al. 2017; Deng et al. 2017; Li et al. 2018; Guo et al. 2018; Fan et al. 2019) (Table 1). Additionally, two QTL for TS were detected on chromosomes 5A and 7A in several studies (Jantasuriyarat et al. 2004; Ding et al. 2011; Wang et al. 2011; Cui et al. 2012; Wu et al. 2012; Xu et al. 2014; Zhai et al. 2016; Ma et al. 2018; Fan et al. 2019) (Table 1).

**Table 1.**
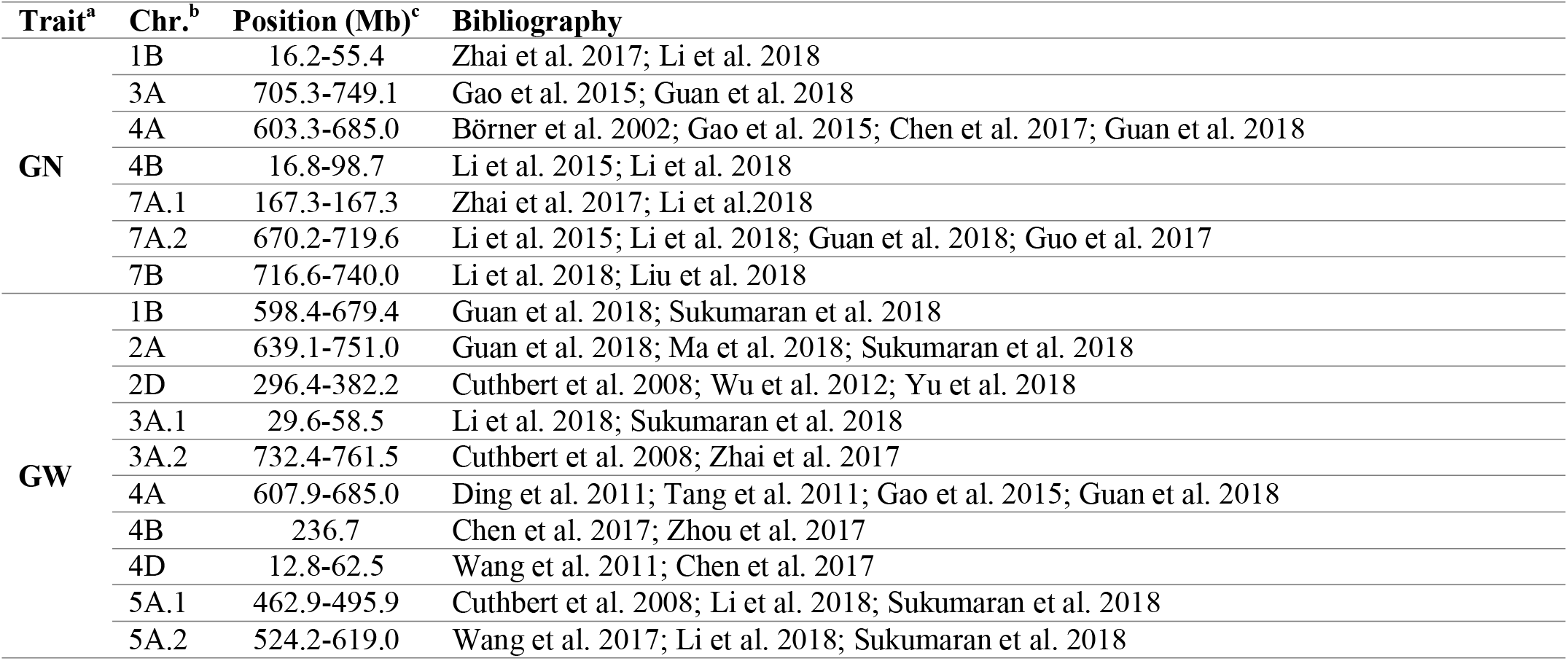
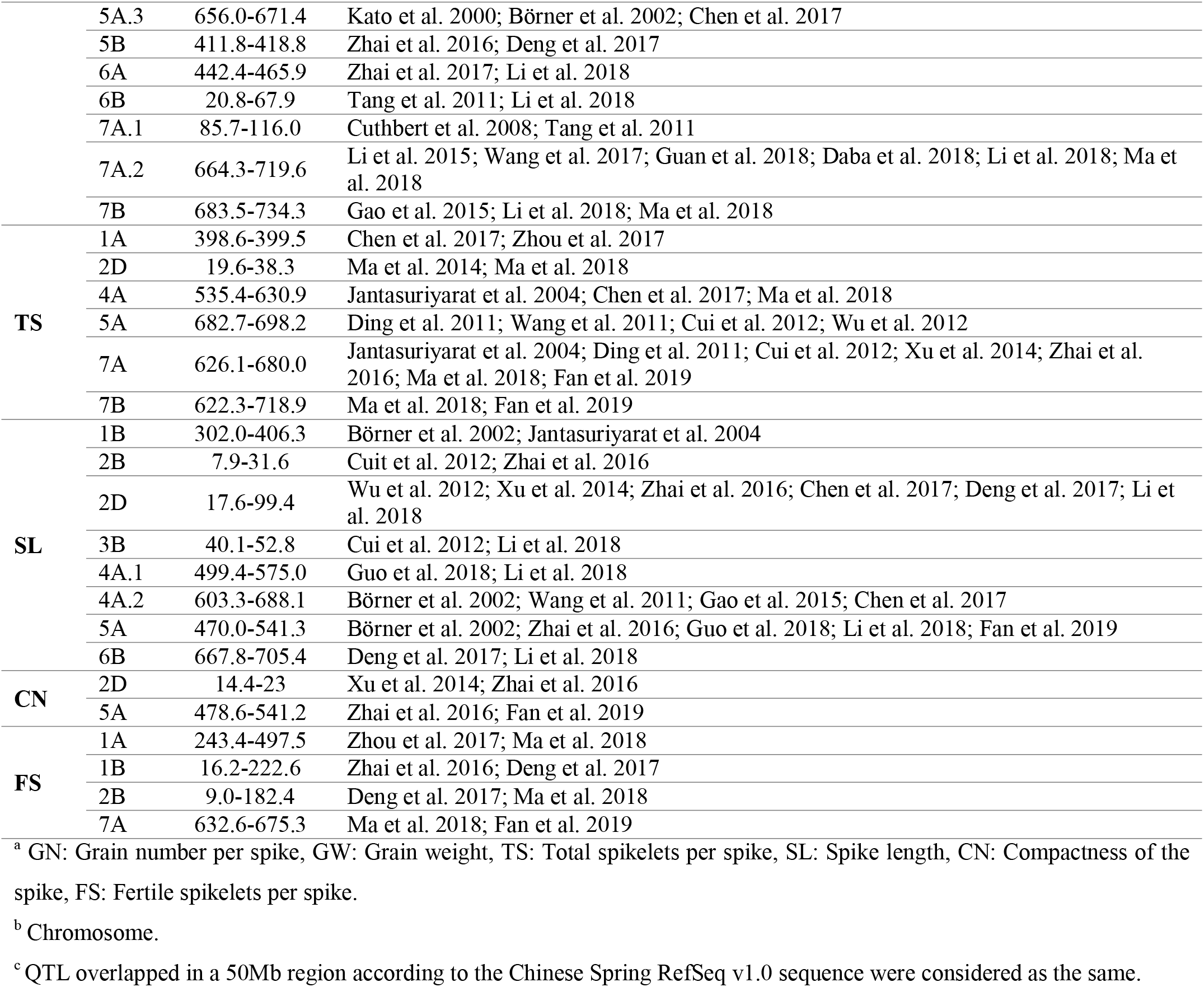
Significant QTL detected in different studies that sheared interval positions according to the Chinese Spring RefSeq v1.0 sequence.

From the crop physiology approach, the GN depends on the florets that reach the fertile stage at anthesis (fertile florets per spike, FF) and on the proportion of them that set grains (grain set, GST, grains per fertile floret). Both depend on the assimilate availability, the first one for the growing spike and developing florets during the 20 days before anthesis (Fischer 1975, 1985; Kirby 1988; González et al. 2011), and the second one during the 10 days after anthesis (Fischer 1975, 1985). This would explain the high importance in GN and FF determination of: (i) the spike dry weight achieved at anthesis (SDW) (Fischer and Stockman 1980; Siddique et al. 1989; Slafer and Andrade 1993; Fischer 1993); and (ii) the dry matter partitioned within the spike between florets/grains and spike structure parts, that is, the fertile floret efficiency (FFE, fertile florets per g of SDW) (Pretini et al. 2020a) and the fruiting efficiency (FE, grains per g of SDW, or FEm grains per g of chaff at maturity) (Abbate et al. 1998; Bustos et al. 2013; García et al. 2014; Elía et al. 2016; Terrile et al. 2017; Lo Valvo et al. 2018; Pretini et al. 2020a). It was recently reported that in modern elite cultivars the SDW was less important to explain GN variation than the fruiting efficiency (Abbate et al. 1998; Rivera-Amado et al. 2019; Bustos et al. 2013; Elía et al. 2016; García et al. 2014; Lo Valvo et al. 2018; Terrile et al. 2017; Pretini et al. 2020a). The GST is considered to be high in relative modern cultivars (i.e., >80% of fertile florets set grains) (Elía et al. 2016; González et al. 2003; Siddique et al. 1989), but recently it was shown it can be as low as 60% (Guo et la. 2017, Pretini et al. 2020a). Then, the amount of assimilates partitioned within the spike to grains or chaff (CH) or its structures (glume, lemma, palea, awns GLPA-, and rachis R-) and the GST and SDW are worthy of study. Only few studies looked for the genetic bases of these traits (Guo et al. 2017; Li et al. 2018, Gerard et al. 2019; Basile et al. 2019; Pretini et al. 2020b, Table S1).

In a previous work (Pretini et al. 2020b), using one of the DH populations use in the present study, we identified and validated the novel *QFEm.perg.3A* for FEm in the chromosome 3A, and the first known QTL for FFE and FE, *QFFE.perg.5A*, located in the chromosome 5A. This last QTL was also detected when the FEm was measured, agreeing with Basile et al. (2019) who detected two regions within this QTL associated to FEm. Despite we studied the pleiotropic effect of both *QFFE.perg.5A* and *QFEm.perg.3A* on the associated traits of spike fertility mentioned previously, we did not look for new major and stable QTL for those traits themselves.

The aim of this study was to identify stable and major QTL for the spike fertility related traits (numerical and physiological components) and to discuss the possible pleiotropic effects among them and with the previously reported *QFEm-perg.3A* and *QFFE-perg.5A*. Two double haploid populations (Baguette Premium 11 x BioINTA 2002 and Baguette 19 x BioINTA2002) derived from the crosses of elite cultivars adapted to the central region of the wheat-producing area of Argentina were used. In the present study we report 305 QTL for different spike fertility related traits distributed throughout the wheat genome. Furthermore, we identify 8 genomic regions that group some of the significant and stable QTL and analyze their pleiotropic effect on other related traits. Finally, we found two regions (R5A1 and R5A.2) and a QTL (*QGW.perg-*6B) that resulted in a final higher yield.

## 2. Materials and methods

### 2.1. Plant materials

Two double haploid (DH) populations were developed from the crosses between Baguette Premium 11 x BioINTA 2002 (BP11xB2002) and Baguette 19 x BioINTA 2002 (B19xB2002). BP11xB2002 consisted of 81 lines whereas B19xB2002 consisted of 102 lines. The three parent lines are semi-dwarf hard hexaploid wheat cultivars and are adapted to central area of wheat production in Argentina (north of Buenos Aires and south of Córdoba provinces). BP11and B19 were released by Nidera Semillas in 2004 and 2006, respectively, in Argentina, while B2002 (BPON/CCTP-F7-7792–122(87)) was developed by CIMMYT (International Maize and Wheat Improvement Center) and released in 2006 in Argentina by INTA. Cycle to anthesis in optimal sowing dates are similar for the three parents (González et al. 2011). The GW of B2002 was higher compared with the GW of BP11 and B19, whereas the GN of B19, followed by BP11, was higher than that of B2002 (Terrile et al. 2017). Generally, B2002 showed higher SDW and chaff than B19 and BP11 (Terrile et al. 2017; Pretini et al. 2020a, b). The three parents are spring cultivars and mostly insensitive to photoperiod (BP11 and B19: *Vrn-A1b/vrn-B1/vrn-D1* and *Ppd-D1a*; B2002: *vrn-A1/Vrn-B1/vrn-D1* and *Ppd-D1a*).

Both populations were genotyped with the iSelect 90K SNP assay (Wang et al. 2014) and evaluated in four (BP11xB19) or five (B19xB2002) environments (E1-E5, Table 2).

**Table 2.**
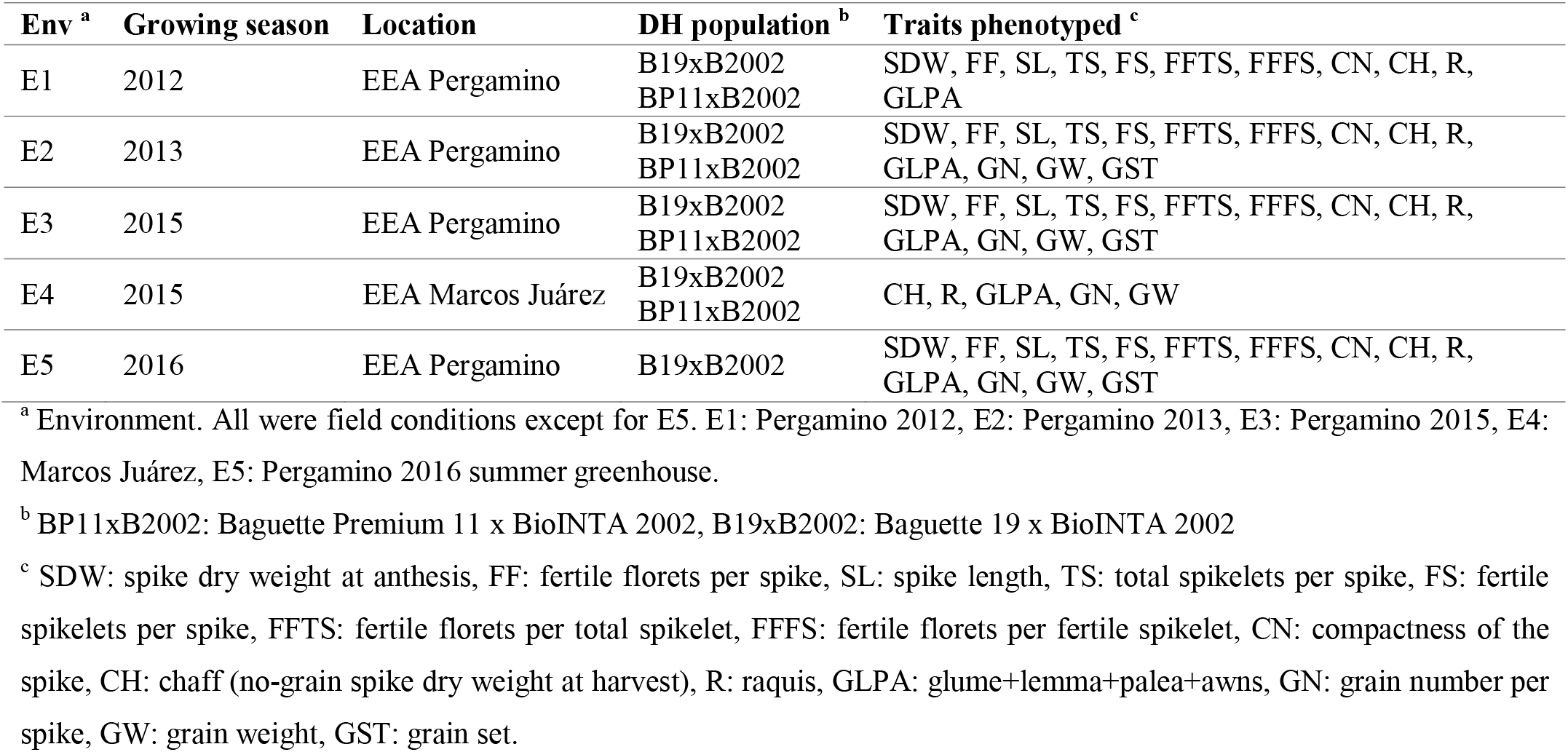
Characteristics of the studied environments. Growing period, location and traits phenotyped for each DH population.

### 2.2. Experiments and Phenotyping

The DH populations were grown in two experimental sites: EEA Pergamino (33° 51’S, 60° 56’W) and EEA Marcos Juárez (32° 43’S, 62° 06’W) Research Stations of INTA (Instituto Nacional de Tecnología Agropecuaria, Argentina) (Table 2). The field trails were carried out during three cropping seasons at Pergamino (E1: 2012, E2: 2013 and E3: 2015) and one cropping season at Marcos Juarez (E4: 2015) (Table 2). The greenhouse trail was carried out only for B19xB2002 during the summer season at Pergamino (E5: 2016) (Table 2). Field experiments were conducted in a randomized complete block design (RCBD) with two replicates. The E1 consisted of double-row plots (1 m long and 0.21 apart) with a sowing rate of 190 plants m^−2^. The E2 to E4 consisted of five-row plots (2 m long and 0.20 apart) with a sowing rate of 330 plants m^−2^ in E2 and 280 plants m^−2^ in E3 and E4. The greenhouse experiment was conducted in a complete randomized design (CRD) with six replicates. Five plants per pot (5 l capacity) were transplanted after being vernalized in cool chamber (20 days at 5°C, 8 h light) during February.

Plants were sampled at the anthesis stage (Z6.1, Zadoks et al. 1974). In E1, five of the most representative spikes of each row were cut. In E2 and E3, a half meter of a central row was sampled, and the spikes were separated from the rest of the biomass, then the spikes were arranged by length and the three median spikes were selected. In E5 two main stem spikes of each pot were cut. In E4 no measurement at anthesis was made. The spike length (SL, mm) was measured from the base of the first spikelet to the terminal spikelet using an electronic caliper. The number of total spikelets per spike was counted (TS) and the spike compactness (CN, mm spikelet node^−1^) was estimated as the ratio between SL and TS. The number of fertile floret (FF) of each spike was count using a binocular microscope. The floret was considered fertile when yellow anthers were visible, or the floret score was >9.5 in Waddington scale (Waddington et al. 1983). The FF was count in a half side of the spike (“a”) and in the terminal spikelet (“t”). The final number was estimated multiplying “a” by two and adding the “t” value. The number of fertile spikelet (FS) was estimated as the number of spikelets with at least one fertile floret multiplied by 2 and adding the terminal spikelet in case it was fertile. The number of fertile florets per total spikelet per spike (FFTS) was estimated as the ratio FF/TS and the number of fertile florets per fertile spikelet per spike (FFFS) as the relation FF/FS. The spike dry weight at anthesis (SDW) was estimated after drying in an oven at 70°C during 48 h.

When plants reached maturity (Z9, Zadoks et al. 1974) a second sample was performed. In E1 to E3 and E5, the spikes were selected following the same criteria as in anthesis. In E4, the spikes were selected following the same criteria as in E2 and E3. All the spikes were dried in an oven at 70°C during 48 h and weighted before threshing by hand. For E1 and E3 to E5, the rachis (R) and the rest of the no-grain parts (glume+lemma+palea+awn, GLPA) were separated when threshing and weighted. For those environments, the chaff (no-grain spike dry weight at harvest) was calculated as the sum of R+GLPA. In contrast, in E2 the chaff was estimated by subtracting the weight of all the grains from the dry weight of the spike before threshing, because no chaff dissection was performed. The grain number (GN) of each spike was counted in E2 to E5 using an automatic counter. The grains from E1 were discard because they were severely affected by fusarium head-blight. The grain weight (GW) was estimated as the ratio between the weight of all grains and the GN. Finally, the grain set (GST) was estimated as de ratio between GN/FF.

### 2.3. Data analyses

For each DH line, the mean value of each trait was calculated across the two replicates for E1 to E4 and the six replicates for E5. The Shapiro–Wilk test and the quantile-quantile (q-q) plot was performed to test for normal distribution. The analysis of variance (ANOVA) was performed using the Infostat/p software (Di Rienzo 2016). In addition, Best Linear Unbiased Estimator (BLUE) was estimated for each DH line including all tested environments as random variable using R v3.3.2.

The narrow-sense heritability of the traits for BP11xB2002 and B19xB2002 was calculated as:

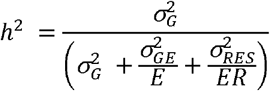

Where σ^2^_G_ is the genotypic (additive) variance, σ^2^_G×E_ is the G×E interaction variance, E is the number of environments, R is the number of replications, and σ^2^_RES_ is the error variance (Hallauer and Miranda 1981).

### 2.4. Linkage map construction and QTL analysis

The DH populations and the three parents were genotyped with the iSelect 90K array containing 90,000 wheat SNP markers (Wang et al. 2014). Additionally, two functional markers for the vernalization genes *Vrn-A1* (Yan et al. 2004) and *Vrn-B1* (Fu et al. 2005) were added to the DH genetic map. Their positions were determined by BLAST against the reference genome IWGSC Ref. Seq. v1.0. (Appels et al. 2018). The SNPs markers with more than 20% of missing and/or heterozygous data were initially discarded for genetic map construction. Then, redundant markers with identical segregations were identified and grouped with the Python script, merger.py^1^. Finally, the R package “Rqtl” (Borman et al. 2003) was used for the genetic map development. The physical position of the SNPs was determined by BLAST against the IWGSC Ref. Seq. v1.0 wheat genome assembly (Appels et al. 2018).

The mean value of the trait in each environment and the BLUE values, which were treated like an additional environment, were used in the QTL mapping. The QTL analysis for the DH populations were scanned with QTL Cartographer 2.5 (Wang 2012) through composite interval mapping (CIM) with the standard model. For the standard model we used a control marker number of 5, a window size of 10 cM and a forward and backward regression method with 500 permutations at α = 0.05. A LOD value of 2.5 was selected as a uniform threshold for all analyses. Detected QTL for a given trait with overlapping support intervals (>50 Mb) were considered as equivalents. The QTL were considered “stable” if they were detected in a minimum of three environments and were defined as “major stable” if they present a R^2^ > 10% in one environment at least.

## 3. Results

### 3.1. Genetic Linkage Map Construction

The linkage map of BP11xB2002 consisted of 7,323 SNPs and two functional markers for the vernalization genes *Vrn-A1* (Yan et al. 2004) and *Vrn-B1* (Fu et al. 2005) (Table S2, S3). All the SNPs represented a total of 723 loci across the 21 wheat chromosomes. The map covered 2605.3 cM in length with an average locus spacing of 4.7 cM (Table S2, S3). The linkage map of B19xB2002 was previously described in Pretini et al. (2020b). Briefly, the B19xB2002 map consisted of 10,936 SNPs and the *Vrn-A1* and *Vrn-B1* markers. All the SNPs represented a total of 739 loci across the 21 wheat chromosomes (Table S4, S5). The map covered 2,221.7 cM in length with an average locus spacing of 4.3 cM (Table S4, S5). Although the genome length of each population was similar, distribution of the markers in the three genomes was not uniform. In BP11xB2002, genomes A and B were similar with at least three times the number of polymorphic markers than genome D, with 2,955, 3,513 and 857 markers, respectively (Table S3). In B19xB2002, the marker uneven distribution was higher with 4,126, 5,448 and 1,364 markers in genomes A, B and D, respectively (Table S5).

The number of *loci* of the genomes A and B triple the number of *loci* of the genome D in both populations, with 311, 300 and 113 *loci* in BP11xB2002 and 324, 317 and 98 *loci* in B19xB2002, respectively (Table S3, S5).

### 3.2. Phenotypic analysis

The means, ranges and heritability of the 13 studied traits across the five environments for the three parents and both DH populations are detailed in Table S6. According to BLUE values, B19 and BP11 parents had higher FF, FFTS, FFFS and GN than B2002, whereas B2002 had higher SDW, SL, TS, CH, R, GLPA and GW (Table 3). BP11 showed the highest FS value whereas B19 showed the lowest FS value and B2002 showed an intermediate FS value (Table 3). All traits showed a normal distribution across each environment and BLUE values, with a transgressive segregation from both parent lines in both populations (Table 3, Table S6).

**Table 3.** Parental and population means, ranges and the Shapiro-Wilk test for all traits based on the adjusted mean (BLUE) values of four (BP11xB2002) and five (B19xB2002) shared environments.

| Trait<sup>a</sup> | Parental Line |  |  | BP11xB2002 |  |  |  |  | B19xB2002 |  |  |  |  |
| --- | --- | --- | --- | --- | --- | --- | --- | --- | --- | --- | --- | --- | --- |
|  | B2002 | BP11 | B19 | Min | Max | Mean | SD<sup>b</sup> | W<sup>c</sup> | Min | Max | Mean | SD | W |
| SL | 98.6 | 93.7 | 87.2 | 86.6 | 122.7 | 101.6 | 7.5 | 0.97 | 79.4 | 105.5 | 94 | 6.4 | 0.96 |
| TS | 22.2 | 19.6 | 20 | 19.6 | 26.4 | 22.7 | 1.6 | 0.96 | 18.7 | 26.1 | 21.1 | 1.4 | 0.98 |
| CN | 4.8 | 4.5 | 4.4 | 3.8 | 5.4 | 4.5 | 0.3 | 0.96 | 3.1 | 5.6 | 4.5 | 0.4 | 0.97 |
| FF | 44.9 | 52.6 | 46.6 | 37.1 | 65.5 | 49.4 | 5.7 | 0.98 | 34.7 | 52.9 | 43.9 | 4.1 | 0.97 |
| FS | 18 | 18.5 | 16.2 | 16.1 | 22.3 | 19 | 1.3 | 0.97 | 14.5 | 19.3 | 17 | 1 | 0.95 |
| FFTS | 2 | 2.4 | 2.3 | 1.7 | 2.7 | 2.2 | 0.2 | 0.95 | 1.3 | 2.5 | 2.1 | 0.2 | 0.96 |
| FFFS | 2.4 | 2.8 | 2.8 | 2.2 | 3.1 | 2.6 | 0.2 | 0.95 | 2 | 3 | 2.5 | 0.2 | 0.96 |
| SDW | 418 | 359 | 357 | 305 | 502 | 408 | 36 | 0.99 | 279 | 480 | 370 | 38 | 0.98 |
| R | 83 | 67 | 61 | 58 | 115 | 79 | 10 | 0.95 | 52 | 94 | 72 | 9 | 0.97 |
| GLPA | 436 | 253 | 346 | 236 | 439 | 335 | 47 | 0.96 | 262 | 478 | 359 | 44 | 0.95 |
| CH | 513 | 315 | 399 | 307 | 535 | 414 | 52 | 0.95 | 323 | 564 | 431 | 50 | 0.96 |
| GN | 37.4 | 40.4 | 39.3 | 29.4 | 53.4 | 39.8 | 5.5 | 0.96 | 27.1 | 50 | 38 | 4.2 | 0.99 |
| GW | 35.8 | 31.1 | 34 | 24.2 | 43.3 | 31.8 | 3.8 | 0.97 | 26.6 | 43.5 | 34.6 | 3.8 | 0.95 |
| GST | 0.88 | 0.83 | 0.87 | 0.59 | 1.11 | 0.87 | 0.1 | 0.98 | 0.5 | 1.3 | 0.9 | 0.2 | 0.92 |

<sup>a</sup> SL: spike length (mm), TS: total spikelets per spike (n° spike<sup>-1</sup>), CN: compactness of the spike (mm node<sup>-1</sup>), FF: fertile florets per spike (n° spike<sup>-1</sup>), FS: fertile spikelets per spike (n° spike<sup>-1</sup>), FFTS: fertile florets per total spikelet (n° spikelet<sup>-1</sup>), FFFS: fertile florets per fertile spikelet (n° spikelet<sup>-1</sup>), SDW: spike dry weight at anthesis (mg spike<sup>-1</sup>), R: raquis (mg spike<sup>-1</sup>), GLPA: glume+lemma+palea+awns (mg spike<sup>-1</sup>), CH: chaff (no-grain spike dry weight at harvest, mg spike<sup>-1</sup>), GN: grain number per spike (n° spike<sup>-1</sup>), GW: grain weight (g), GST: grain set.
<sup>b</sup> SD: standard error
<sup>c</sup> W: A modification of the test of Shapiro-Wilks for normality. Mahibbur and Govindarajulu (1997).

The phenotypic performances of the GN, FF, SDW, GST and CH in both populations were already described in Pretini et al. (2020a). Briefly, the GN ranged from 29.4 to 53.4 and from 27.2 to 50.0 grains per spike in the BP11xB2002 and B19xB2002 populations, respectively. The FF ranged from 37.1 to 65.5 and from 34.7 to 52.9 florets per spike in the BP11xB2002 and B19xB2002 populations, respectively. The SDW ranged from 304 to 502 and from 279 to 480 mg per spike in the BP11xB2002 and B19xB2002 populations. The GST ranged from 0.59 to 1.11 and from 0.5 to 1.3 in the BP11xB2002 and B19xB2002 populations, respectively (Table S6).

In relation to the traits determining spike structure at anthesis, the highest SL, considering the agronomic environments (E1 to E4) and both populations, was observed in E1 (109.5 ± 10.6 and 105.8 ± 10.7 mm, for BP11xB2002 and B19×2002, respectively), while the lowest SL was observed in E2 (93.6 ± 7.4 and 92.5 ± 6.9 mm, for BP11xB2002 and B19×2002, respectively) (Table S6). The E5 showed even a lower SL than E2 with 73.4 ± 8.4 mm (Table S6). The TS ranges for both DH populations were similar within the agronomic environments (from 21.3 ± 1.6 to 23.9 ± 1.9 spikelets per spike for BP11xB2002 and from 21.0 ± 1.3 to 23.7 ± 2.0 spikelets per spike for B19xB2002, E2 to E1 in both cases); while the lowest TS was observed in the non-agronomic environment (E5, 16.2 ± 2.4 spikelets per spike, Table S6). Also, the FS ranges were similar between both populations (from 17.5 ± 1.5 to 20.6 ± 1.6 fertile spikelets per spike for BP11×2002 and from 17.3 ± 1.2 to 20.1 ± 1.6 fertile spikelets per spike for B19xB2002, E2 to E1 in both cases) and the lowest FS was observed in the E5 (11.5 ± 1.4 fertile spikelets per spike, Table S6). The lowest CN for both populations was detected in the E2 (4.4 ± 0.3 mm per spikelet, Table S6). In contrast, the highest CN varied according to the population × environment combination (for BP11xB2002 it was detected in E1 with 4.6 ± 0.4 mm per spikelet, whereas for B19xB2002 it was detected in the E5 with 4.6 ± 0.5 mm per spikelet, Table S6). In general, for both populations, as the spikes were longer (higher SL), the TS increased, and CN decreased (Table S6).

The FFTS ranged from 2.0 ± 0.2 to 2.3 ± 0.4 fertile florets per fertile spikelet for BP11xB2002 and from 2.1 ± 0.3 to 2.4 ± 0.4 fertile florets per fertile spikelet for B19xB2002 (E2 to E1, Table S6). However, the lowest FFTS was detected in the E5 with 1.7 ± 0.3 fertile florets per fertile spikelet (Table S6). The FFFS ranged from 2.4±0.2 to 2.7±0.2 (E5 to E3) fertile florets per fertile spikelet for BP11xB2002 and from 2.40±0.2 to 2.6±0.3 (E2 to E1) fertile florets per fertile spikelet for B19xB2002 (Table S6).

As regards to the CH at maturity, for both populations within the agronomic environment, it was the highest in E3 (598 ± 132 mg per spike for BP11xB2002 and 663 ± 104 mg per spike for B19xB2002) and the lowest in E2 (277 ± 41 mg per spike for BP11xB2002 and 277 ± 41 mg per spike in B19xB2002). The non-agronomic environment E5 reached a similar value to E2 (282 ± 80 mg per spike). The spike dry matter partitioning at maturity between R and GLPA varied from 14 to 22% for the R, and from 78 to 86% for the GLPA, depending on DH population and environment. Similarly to CH within the agronomic environments, the highest R was detected in the E3 (97 ± 19 mg for BP11xB2002 and 100 ± 16 mg for B19xB2002), and the lowest in the E2 (60 ± 10 mg per spike for BP11xB2002 and 59 ± 7 mg per spike for B19xB2002, Table S6). The R detected in the E5 was even lower than the one of E2 (48 ± 13 mg per spike, Table S6). The GLPA ranged in the agronomic environments E3 to E2 from 217 ± 35 to 502 ± 116 mg per spike for BP11xB2002 and from 215 ± 39 to 564 ± 93 mg per spike for B19xB2002 (Table S6), being 234 ± 70 mg per spike in E5. The GW ranged from 31.5 ± 4.7 to 32.6 ± 5.0 g (E4-E3) for BP11xB2002 and from 29.2 ± 3.4 to 40.6 ± 6.1 g (E2-E5) for B19xB2002 including the non-agronomic trait (Table S6).

In BP11xB2002, the h^2^ ranged from 0.31 to 0.86, with the lowest value in the SDW and the highest value in TS (Table S6). Meanwhile, in B19xB2002, the h^2^ ranged from 0.36 to 0.68, with the lowest value also in SDW and the highest in FFFS (Table S6).

### 3.3. QTL Mapping Analysis

A total of 305 QTL were identified across 5 environments and BLUE distributed on the 21 chromosomes (Table S7). However, only 28 QTL were present in at least 3 individual environments or BLUE analysis with a LOD > 2.5 considering a single population or a combination of both populations but with the contribution of the same germplasm (Baguette or B2002) and were considered stable with R^2^ > 10% in one environment at least (Table 4). Those stable QTL were distributed on the 1A, 2A, 2B, 2D, 3A, 3B, 5A, 5B, 6A, 6B, 7A and 7B chromosomes (Table 4).

**Table 4.**
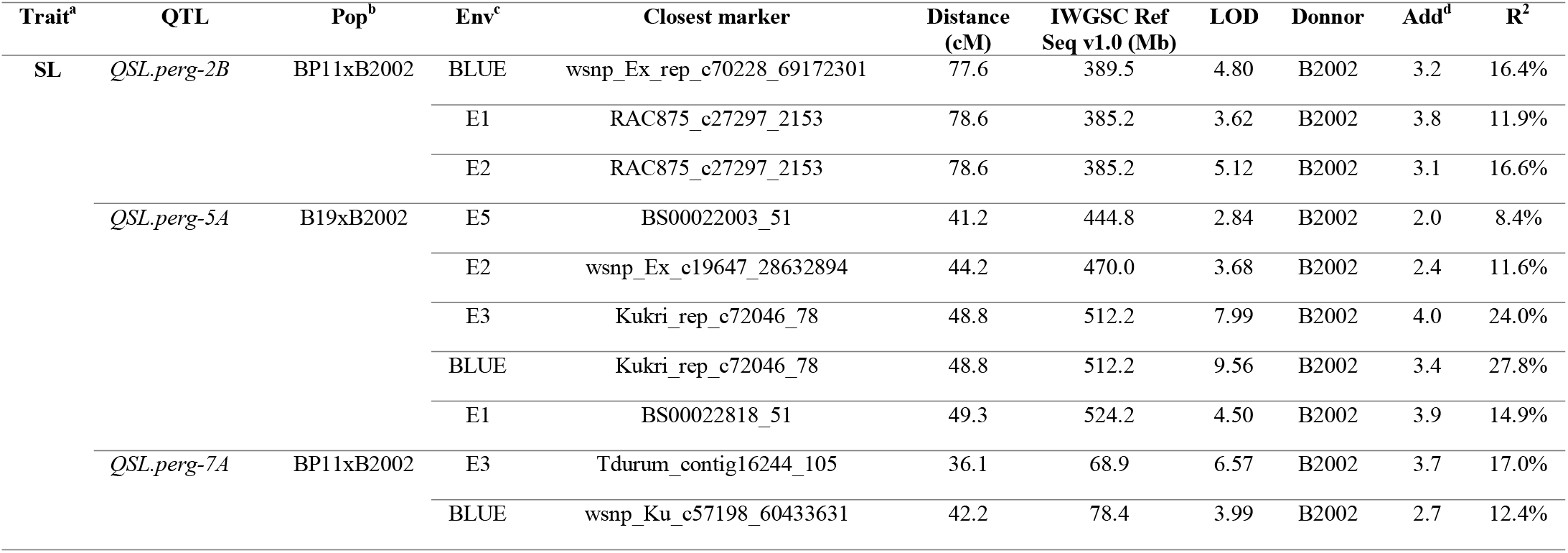
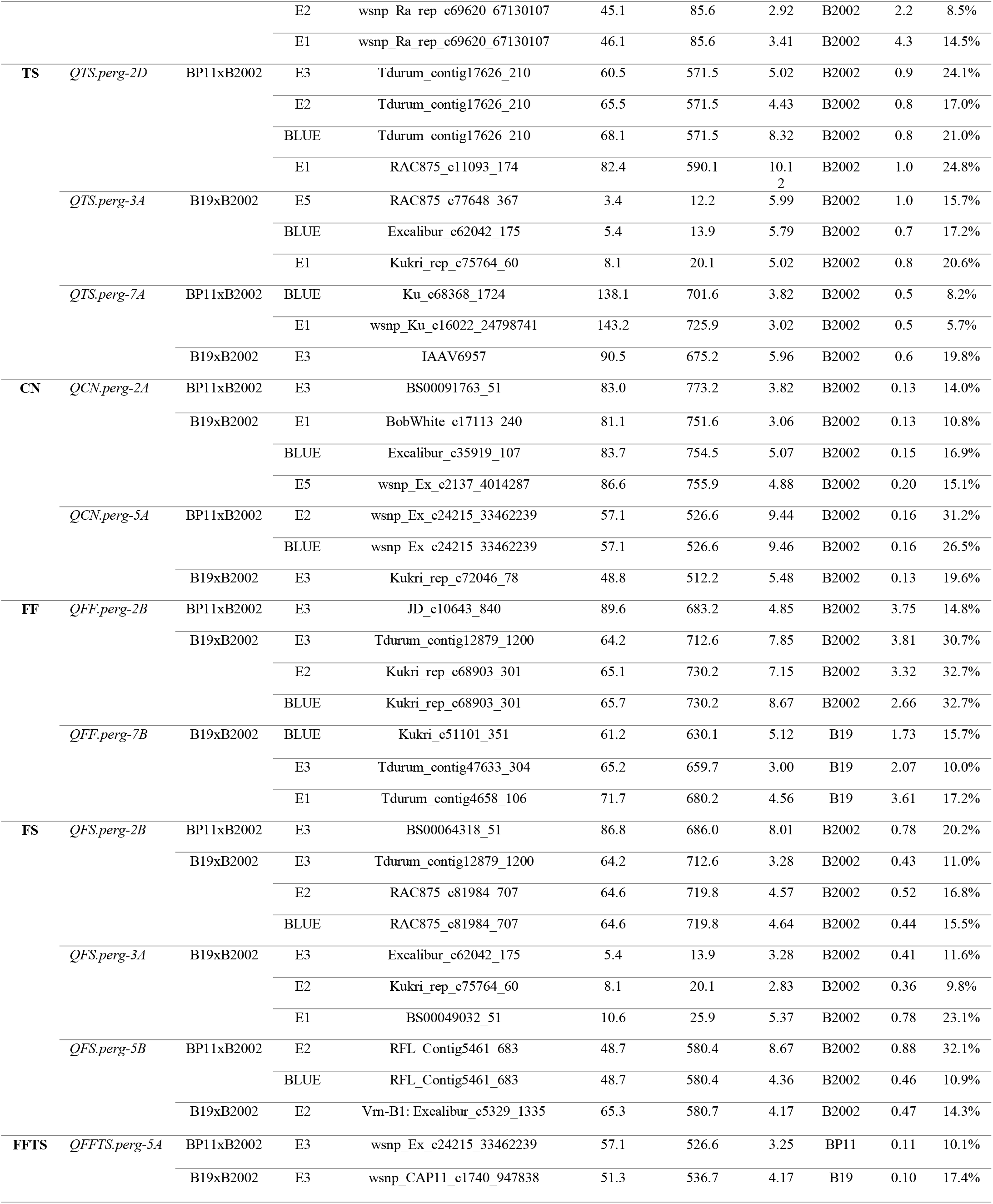
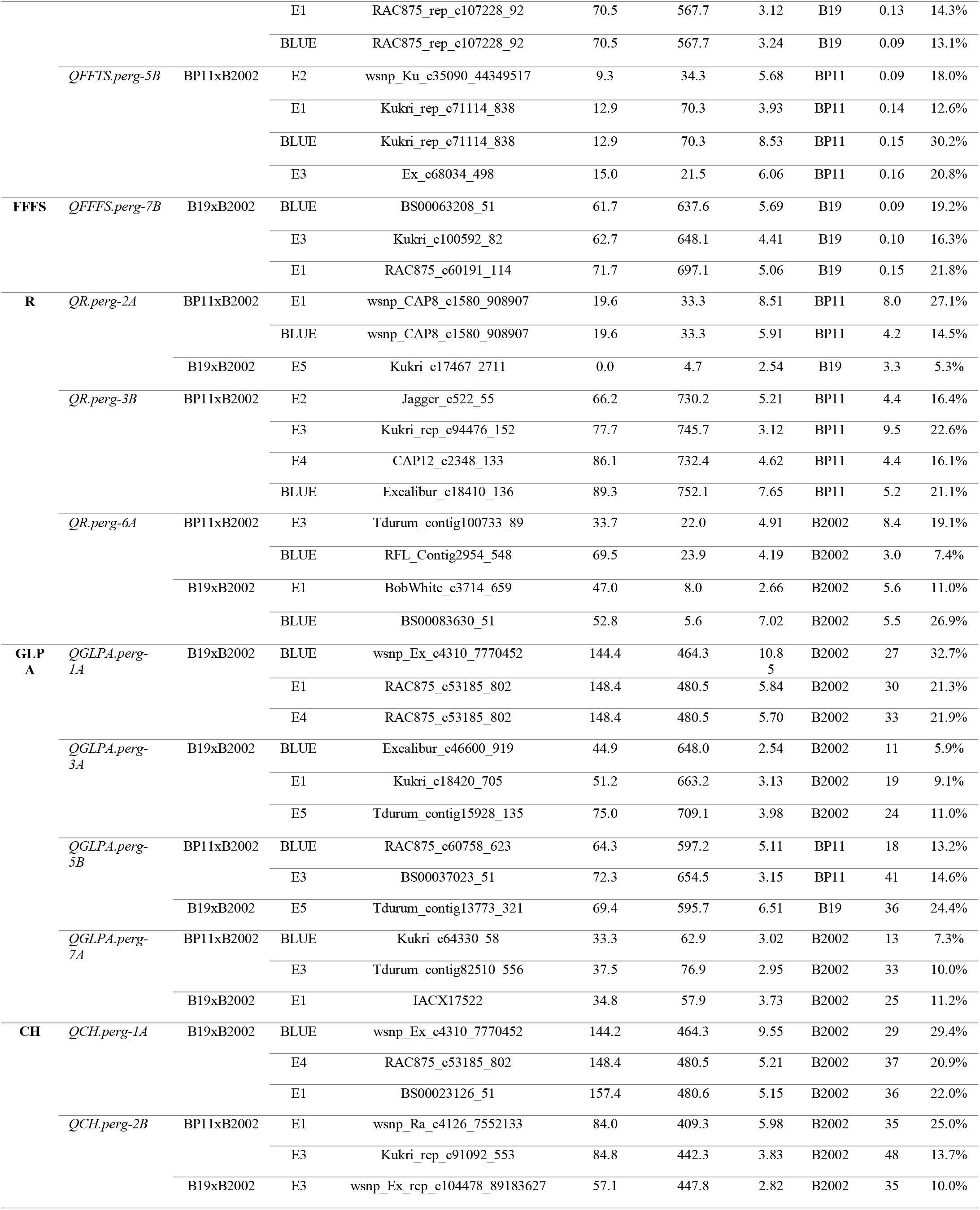
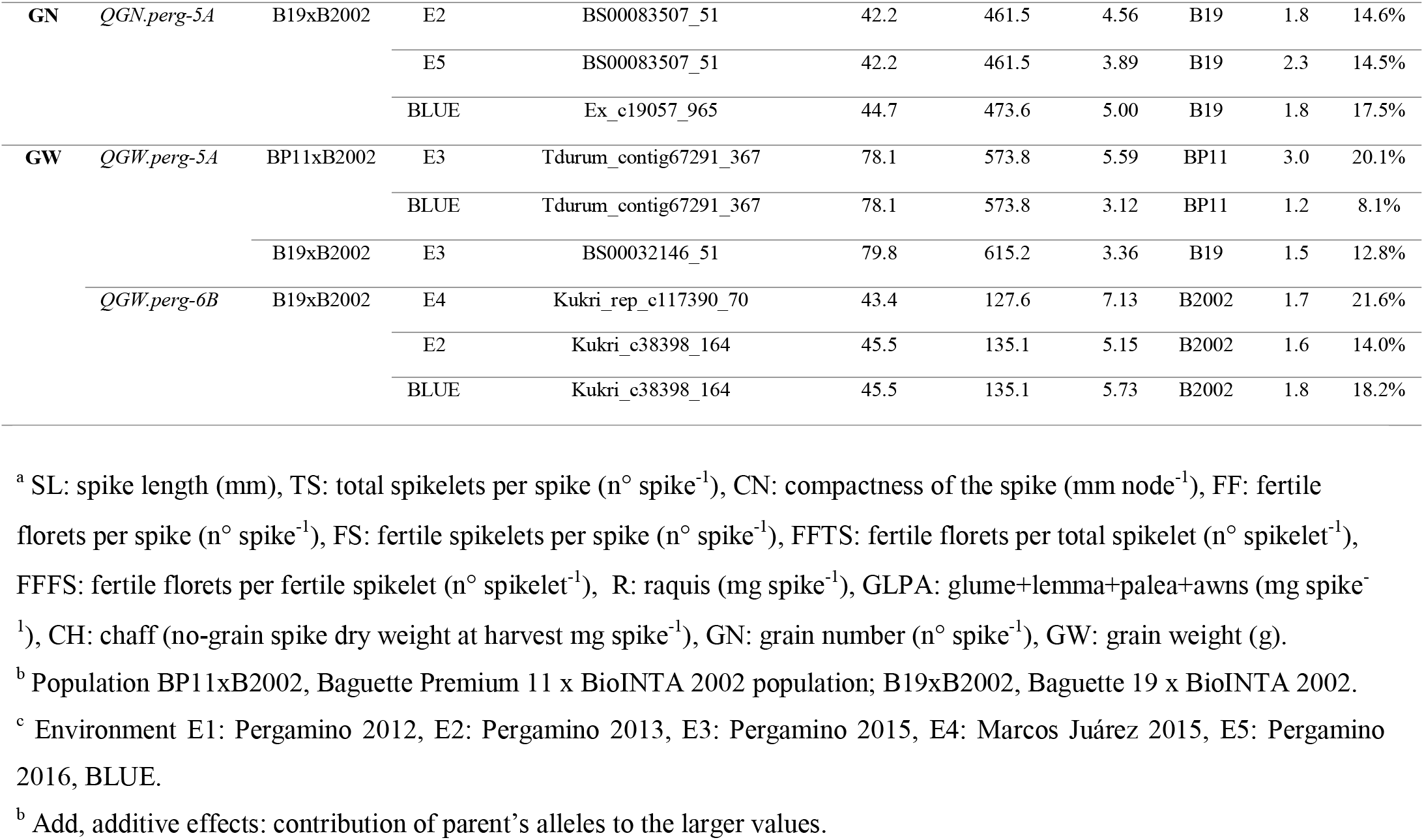
Stable and major QTL identified for spike fertility related traits in both populations.

#### 3.3.1. QTL for SL

The QTL analysis identified 19 regions for SL on chromosomes 1A, 1B, 2A, 2B, 2D, 3B, 4B, 5A, 5B, 6A, 6D and 7A with a LOD score that ranged from 2.54 to 9.56, explaining 4.7-27.8% of the phenotypic variance (Table S7). However, three QTL on chromosomes 2B (*QSL.perg-2B*), 5A (*QSL.perg-5A*) and 7A (*QSL.perg-7A*) were consider major and stable across environments. Both QTL detected for *QSL.perg-2B* and *QSL.perg-7A* were from the BP11xB2002 population while the *QSL.perg-5A* was detected in the B19xB2002 population (Table 4).

The QTL peak of *QSL.perg-2B* was mapped at the RAC875_c27297_2153 SNP marker (78.6 cM, 385.2 Mb) with a LOD of 5.12 and a R^2^ of 16.6% (Table 4). The QTL peak of *QSL.perg-5A* was mapped at the Kukri_rep_c72046_78 SNP marker (48.8 cM, 512.2 Mb) with a LOD of 9.56 and a R^2^ of 27.8% (Table 4). Finally, the QTL peak of *QSL.perg-7A* was mapped at the Tdurum_contig16244_105 SNP marker (36.1 cM, 68.9 Mb) with a LOD of 6.57 and a R^2^ of 17.0% (Table 4). The increasing allele was always contributed by B2002 with an additive effect that ranged from 3.1 to 3.8 mm for *QSL.perg-2B*, from 2.0 to 4.0 mm for *QSL.perg-5A* and from 2.2 to 4.3 mm for *QSL.perg-7A* (Table 4).

#### 3.3.2. QTL for TS

The QTL analysis identified 24 regions for TS on chromosomes 1B, 1D, 2A, 2B, 2D, 3A, 4B, 5A, 5B, 5D, 6A, 6B, 7A and 7B with a LOD score that ranged from 2.77 to 10.12, explaining 5.7-28.2% of the phenotypic variance (Table S7). However, three QTL on chromosomes 2D (*QTS.perg-2D*), 3A (*QTS.perg-3A*) and 7A (*QTS.perg-7A*) were considered major and stable across environments. The *QTS.perg-2D* was detected in the BP11xB2002 population while the *QTS.perg-3A* was observed in the B19xB2002 population (Table 4). The *QTS.perg-7A* was detected in two environments in the BP11xB2002 population (E1 and BLUE), and in one environment if the B19xB2002 population (E3, Table 4).

The QTL peak of *QTS.perg-2D* was mapped at the RAC875_c11093_174 SNP marker (82.4 cM, 590.1 Mb) with a LOD of 10.12 and R^2^ of 24.8% (Table 4). The QTL peak of *QTS.perg-3A* was mapped at the RAC875_c77648_367 SNP marker (3.4 cM, 12.2 Mb) with a LOD of 5.99 and R^2^ of 15.7% (Table 4). Finally, the QTL peak of *QTS.perg-7A* was mapped at the IAAV6957 SNP marker (90.5 cM, 675.2 Mb) with a LOD of 5.96 and a R^2^ of 19.8% (Table 4). The increasing allele for all the QTL were contributed by B2002 with an additive effect that ranged from 0.8 to 1.0 total spikelets per spike for *QTS.perg-2D*, from 0.7 to 1.0 total spikelets per spike for *QTS.perg-3A* and from 0.5 to 0.6 total spikelets per spike for *QTS.perg-7A* (Table 4).

#### 3.3.3. QTL for CN

The QTL analysis identified 22 QTL for CN on chromosomes 1B, 1D, 2A, 3A, 3B, 4B, 5A, 5B, 6A, 6D, 7A and 7B with a LOD score that ranged from 2.55 to 9.46, explaining 6.7-31.2% of the phenotypic variance (Table S7). However, two QTL on chromosomes 2A (*QCN.perg-2A*) and 5A (*QCN.perg-5A*) were considered major and stable across environments. The QTL were identified in both populations. The *QCN.perg-2A* was detected in one environment in BP11xB2002 (E3) and in three environments in B19xB2002 (E1, E5 and BLUE). The *QCN.perg-5A* was detected in two environments in BP11xB2002 (E2 and BLUE) and in one environment in B19xB2002 (E3, Table 4).

The peak of *QCN.perg-2A* was mapped at the Excalibur_c35919_107 SNP marker (83.7 cM, 754.5 Mb) with a LOD of 5.07 and a R^2^ of 16.9% (Table 4). The peak of *QCN.perg-5A* was mapped at the wsnp_Ex_c24215_33462239 SNP marker (57.1 cM, 526.6 Mb) with a LOD of 9.46 and R^2^ of 26.5% (Table 4). The increasing allele for both QTL was contributed by B2002 and had an additive effect ranged from 0.13 to 0.20 mm per node (Table 4).

#### 3.3.4. QTL for FF

The QTL analysis identified 18 regions for FF on chromosomes 1A, 2A, 2B, 3A, 3B, 3D, 4B, 5A, 5B, 5D, 6A and 7B. The LOD score ranged from 2.54 to 7.85, explaining 5.5-32.7% of the phenotypic variation (Table S7). However, two QTL on chromosomes 2B (*QFF.perg-2B*) and 7B (*QFF.perg-7B*) were consider major and stable across environments. *QFF.perg-2B* was detected in one environment in BP11xB2002 (E3) and three environments in B19xB2002 (E2, E3 and BLUE, Table 4). While the *QFF.perg-7B* was detected in three environments of the B19xB2002 population (Table 4).

The QTL peak of *QFF.perg-2B* was mapped at the Kukri_rep_c68903_301 SNP marker (65.7 cM, 730.2 Mb) with a LOD of 8.67 and a R^2^ of 32.7% (Table 4). The increasing allele of *QFF.perg-2B* was contributed by B2002 and had a significant additive effect that ranged from 3.3 to 3.8 fertile florets per spike. The QTL peak of *QFF.perg-7B* was mapped at the Kukri_c51101_351 SNP marker (61.2 cM, 630.1 Mb) with a LOD of 5.12 and a R^2^ of 15.7% (Table 4). The increasing allele of *QFF.perg-7B* was contributed by the B19 and had a significant additive effect that ranged from 1.7-3.6 fertile florets per spike (Table 4).

#### 3.3.5. QTL for FS

The QTL analysis identify 23 regions for FS on chromosomes 1D, 2A, 2B, 3A, 3B, 3D, 4A, 5A, 5B, 5D, 6A, 6B, 7A and 7B with a LOD score that ranged from 2.55 to 8.67, explaining 6.2-32.1% of the phenotypic variance (Table S7). However, three QTL on chromosomes 2B (*QFS.perg-2B*), 3A (*QFS.perg-3A*) and 5B (*QFS.perg-5B*) were considered major and stable across environments. The *QFS.perg-2B* was detected in one environment in B11xB2002 (E3) and in three environments in B19xB2002 (E2, E3 and BLUE, Table 4). The *QFS.perg-3A* was detected in three environments in the B19xB2002 population (E1, E2 and E3) (Table 4). Finally, the *QFS.perg-5B* was detected in two environments in BP11xB2002 (E2 and BLUE) and in one environment in B19xB2002 (E2, Table 4).

The QTL peak of *QFS.perg-2B* was mapped at the BS00064318_51 SNP marker (86.8 cM, 686.0 Mb) with a LOD of 8.01 and a R^2^ of 20.2% (Table 4). The QTL peak of *QFS.perg-3A* was mapped at the BS00049032_51 SNP marker (10.6 cM, 25.9 Mb) with a LOD of 5.37 and a R^2^ of 23.1% (Table 4). Finally, the QTL peak of *QFS.perg-5B* was mapped at the RFL_Contig5461_683 SNP marker (48.7 cM, 580.4 Mb) with a LOD of 8.67 and a R^2^ of 32.1% (Table 4). The increasing allele was always contributed by B2002 with an additive effect that ranged from 0.4 to 0.9 fertile spikelets per spike (Table 4).

#### 3.3.6. QTL for FFTS

The QTL analysis identified 21 regions for FFTS on the chromosomes 1B, 2A, 2B, 2D, 3A, 3B, 3D, 4B, 5A, 5B, 6A, 7A, 7B and 7D with a LOD score that ranged from 2.52 to 8.53, explaining 7.3-30.8% of the phenotypic variance (Table S7). However, two QTL on chromosomes 5A (*QFFTS.perg-5A*) and 5B (*QFFTS.perg-5B*) were considered major and stable across environments. The *QFFTS.perg-5A* was detected in one environment in BP11xB2002 (E3) and in three environments in B19xB2002 (E1, E3 and BLUE) while the *QFFTS.perg-5B* was were detected in four environments in BP11xB2002 (E1, E2, E3 and BLUE) (Table 4).

The QTL peak for *QFFTS.perg-5A* was mapped at the wsnp_CAP11_c1740_947838 SNP (51.3 cM, 536.7 Mb) with a LOD of 4.17 and a R^2^ of 17.4% (Table 4). The QTL peak for *QFFTS.perg-5B* was mapped at the Kukri_rep_c71114_838 SNP (12.9 cM, 70.3 Mb) with a LOD of 8.53 and a R^2^ of 30.2% (Table 4). For both QTL, the increasing allele was contributed by the Baguette parents with an additive effect that ranged from 0.09 to 0.13 and from 0.9 to 0.16 fertile florets per total spikelet per spike for *QFFTS.perg-5A* and *QFFTS.perg-5B*, respectively (Table 4).

#### 3.3.7. QTL for FFFS

The QTL analysis identified 22 QTL for FFFS on chromosomes 1A, 1B, 2A, 2B, 3A, 3B, 4A, 4B, 5A, 5B, 6B, 7A, 7B and 7D with a LOD score that ranged from 2.58 to 7.65, explaining 6.8-25.4% of the phenotypic variance (Table S7). However, only one QTL on chromosome 7B (*QFFFS.perg-7B*) was considered major and stable across environments. This QTL was detected in the B19xB2002 population (Table 4).

The peak of *QFFFS.perg-7B* was mapped at the BS00063208_51 (61.7 cM, 637.3 Mb) with a LOD of 5.69 and a R^2^ of 19.2% (Table 4). The increasing allele was contributed by B19 with an additive effect that ranged from 0.09 to 0.15 fertile florets per fertile spikelet per spike (Table 4).

#### 3.3.8. QTL for SDW

The QTL analysis identified 24 regions for SDW on chromosomes 1A, 1B, 2B, 2D, 3B, 3D, 4A, 4B, 4D, 5A, 5B, 7A, 7B and 7D with a LOD score that ranged from 2.52 to 7.20 explaining 0.4-37.3% of the phenotypic variance (Table S7). However, no QTL was considered major and stable across environments.

#### 3.3.9. QTL for R

The QTL analysis identified 31 QTL for R on chromosomes 1A, 2A, 2B, 2D, 3A, 3B, 3D, 4A, 4B, 4D, 5A, 5B, 6A, 6B, 6D and 7B with a LOD score that ranged from 2.54 to 8.76, explaining 5.1-30.2% of the phenotypic variance (Table S7). However, three QTL on chromosomes 2A (*QR.perg-2A*), 3B (*QR.perg-3B)* and 6A (*QR.perg-6A*) were considered major and stable across environments. The *QR.perg-3B* was only detected in BP11xB2002 while the *QR.perg-2A* and *QR.perg-6A* were detected in both populations. The *QR.perg-2A* was detected in two environments in BP11xB2002 (E1, BLUE) and in one environment in B19xB2002 (E5) (Table 4). While the *QR.perg-6A* was detected in two environments for each DH population (Table 4).

The peak of *QR.perg-2A* was mapped at the wsnp_CAP8_c1580_908907 SNP marker (19.6 cM, 33.3 Mb) with a LOD of 8.51 and R^2^ of 27.1% (Table 4). The peak of *QR.perg-3B* was mapped at the Ku Excalibur_c18410_136 SNP marker (89.3 cM, 752.1 Mb) with a LOD of 7.65 and R^2^ of 21.2% (Table 4). Finally, the peak of *QR.perg-6A* was mapped at the BS00083630_51 SNP marker (52.8 cM, 5.6 Mb) with a LOD of 7.02 and a R^2^ of 26.9% (Table 4). The increasing allele of *QR.perg-2A* and *Qr. perg-3B* was contributed by the Baguette parents with an additive effect that ranged from 42 to 80 and from 44 to 95 mg per spike, respectively. In contrast, the increasing allele of *QR.perg-6A* was contributed by B2002 with an additive effect that ranged from 30 to 84 mg per spike (Table 4).

#### 3.3.10. QTL for GLPA

The QTL analysis identified 23 QTL for GLPA on chromosomes 1A, 1B, 1D, 2B, 2D, 3A, 3B, 4B, 5A, 5B, 5D, 6A, 7A and 7B with a LOD score that ranged from 2.53 to 10.85, explaining 5.6-32.7% of the phenotypic variance (Table S7). However, four QTL on chromosomes 1A (*QGLPA.perg-1A*), 3A (*QGLPA.perg-3A*), 5B (*QGPLA.perg-5B*) and 7A (*QGLPA.perg-7A*) were considered major and stable across environments. The *QGLPA.perg-1A* and *QGLPA.perg-3A* were detected in BP19xB2002, while the *QGPLA.perg-5B* and *QGLPA.perg-7A* were detected in both populations. The *QGPLA.perg-5B* and *QGLPA.perg-7A* were detected in two environments in BP11xB2002 and in one environment in B19xB2002 (Table 4).

The peak of *QGLPA.perg-1A* was mapped at the wsnp_Ex_c4310_7770452 SNP marker (144.4 cM, 464.3 Mb) with a LOD of 10.85 and a R^2^ of 32.2% (Table 4). The peak of *QGLPA.perg-3A* was mapped at the Tdurum_contig15928_135 SNP marker (75.0 cM, 709.1 Mb) with a LOD of 3.98 and a R^2^ of 11.0% (Table 4). The peak of *QGPLA.perg-5B* was mapped at the Tdurum_contig13773_321 (69.4 cM, 595.7 Mb) with a LOD of 6.52 and a R^2^ of 24.4% (Table 4). Finally, the peak of *QGLPA.perg-7A* was mapped at the IACX17522 SNP marker (34.8 cM, 57.9 Mb) with a LOD of 3.73 and a R^2^ of 11.2% (Table 4). The increasing allele for *QGLPA.perg-1A*, *QGLPA.perg-3A* and *QGLPA.perg-7A* was contributed by B2002, with an additive effect that ranged from 27 to 33, from 11 to 24 and from 13 to 33 mg per spike, respectively (Table 4). On the other and, the increasing allele for *QGLPA.perg-5B* was contributed by the Baguette parent with an additive effect that ranged from 18 to 41 mg per spike (Table 4).

#### 3.3.11. QTL for CH

The QTL analysis identified 25 QTL for CH on chromosomes 1A, 1B, 1D, 2A, 2B, 2D, 3A, 3B, 5A, 5B, 6A, 7A, 7B and 7D with a LOD score that ranged from 2.53 to 9.55, explaining 6.4-35.9% of the phenotypic variance (Table S7). However, two QTL on chromosomes 1A (*QCH.perg-1A*) and 2B (*QCH.perg-2B*) were considered major and stable across environments. The *QCH.perg-1A* was detected in B19xB2002 while the *QCH.perg-2B* was detected in two environments in B11xB2002 (E1 and E3) and in one environment in B19xB2002 (E3) (Table 4).

The peak of *QCH.perg-1A* was mapped at the wsnp_Ex_c4310_7770452 SNP marker (144.2 cM, 464.3 Mb) with a LOD of 9.55 and a R^2^ of 29.4% (Table 4). The peak of *QCH.perg-2B* was mapped at the wsnp_Ra_c4126_7552133 SNP marker (84.0 cM, 409.3 Mb) with a LOD of 5.98 and a R^2^ of 25% (Table 4). In both cases, the increasing allele was contributed by B2002, with an additive effect that ranged from 29 to 37 mg for *QCH.perg-1A* and from 35 to 48 mg for *QCH.perg-2B* (Table 4).

#### 3.3.12. QTL for GN

The QTL analysis identified 18 QTL for GN on chromosomes 1D, 2A, 2B, 3D, 4D, 5A, 5B, 5D, 6A, 6D, 7B and 7D with a LOD score that ranged from 2.52 to 6.60, explaining 8.0-35.0% of the phenotypic variance (Table S7). However, only one QTL on chromosome 5A (*QGN.perg-5A*) was considered major and stable across environments. In this case, the QTL was detected in B19xB2002 (Table 4).

The peak of *QGN.perg-5A* was mapped at the Ex_c19057_965 SNP marker (44.7 cM, 473.6 Mb) with a LOD of 5.0 and a R^2^ of 17.5% (Table 4). The increasing allele for *QGN.perg-5A* was contributed by B19 with an additive effect that ranged from 1.8 to 2.3 grains per spike (Table 4).

#### 3.3.13. QTL for GW

The QTL analysis identified 21 QTL for GW on chromosomes 1A, 1B, 2A, 2B, 3A, 3D, 5A, 5B, 6A, 6B, 7A, 7B and 7D with a LOD score that ranged from 2.55 to 7.13, explaining 6.8-23.3% of the phenotypic variance (Table S7). However, two QTL on chromosomes 5A (*QGW.perg-5A*) and 6B (*QGW.perg-6B*) were considered major and stable across environments. The *QGW.perg-6B* was detected in B19XB2002 while the *QGW.perg-*5A was detected in tow environments in BP11xB2002 (E3 and BLUE) and in one environment in B19xB2002 (Table 4).

The peak of *QGW.perg-5A* was mapped at the Tdurum_contig67291_367 SNP marker (78.1 cM, 573.8 Mb) with a LOD of 5.59 and a R^2^ of 20.2% (Table 4). Finally, the peak of *QGW.perg-6B* was mapped at the Kukri_rep_c117390_70 SNP marker (43.4 cM, 127.6 Mb) with a LOD of 7.13 and a R^2^ of 21.6% (Table 4). Baguette parents with an additive effect that ranged from 1.2 to 3.0 g contributed to the increasing allele for *QGW.perg-5A* (Table 4). For *QGW.perg-6B*, the increasing allele was contributed by B2002 with an additive effect that ranged from 1.6 to 1.8 g (Table 4).

#### 3.3.14. QTL for GST

The QTL analysis identified 14 QTL for GST on chromosomes 1D, 2B, 2D, 3A, 3B, 4B, 5A, 5B, 5D, 6A and 7B with a LOD score that ranged from 2.54 to 4.95, explaining 9.0-20.1% of the phenotypic variance (Table S7). However, no QTL was considered major and stable across environments.

### 3.4. Stable and major QTL regions for spike fertility related traits

A total of 8 genomic regions distributed in 7 chromosomes (R1A, R2B, R3A, R5A.1, R5A.2, R5B, R7A and R7B) were identified containing 17 of the 28 stable and major QTL detected for the different traits (Table 5). The QTL located in these regions shared a confident interval of +/− 50 Mb from the SNP marker with the highest LOD value according to their physical position, indicating a potential pleiotropic effect on the corresponding traits (Table 5). The increasing alleles for R1A, R2B, R3A and R7A were always contributed by B2002. The QTL peak of the R1A region was located between 464.3-480.6 Mb (+/− 1 LOD) and harbored *QCH.perg-1A* and *QGLPA.perg-1A*, the QTL peak of the R2B region was located between 544.8-741.9 Mb (+/− 1 LOD) and harbored *QFF.perg-2B* and *QFS.perg-2B*, the QTL peak of the R3A region was located between 1.9-32.1 Mb (+/− 1 LOD) and harbored *QTS.perg-3A* and *QFS.perg-3A*, and the QTL peak of the R7A region was located between 36.9-120.2 Mb (+/− 1 LOD) and harbored *QSL.perg-7A* and *QGLPA.perg-7A* (Table 5). On the other hand, the Baguette parents contributed the increasing alleles for the R5A.2 and R7B regions. The QTL peak of the R5A.2 region was located between 470.0-637.5 Mb (+/− 1 LOD) and harbored *QFFTS.perg-5A* and *QGW.perg-5A* and, the QTL peak of the R7B region was located between 605.4-709.3 Mb (+/− 1 LOD) and harbored *QFF.perg-7B* and *QFFFS.perg-7B* (Table 5). Different parents depending on the trait contributed the increasing allele for R5A.1 (Table 5). The QTL peak of the R5A.1 region was located between 389.7-540.6 Mb (+/− 1 LOD) and harbored *QSL.perg-5A* and *QCN.perg-5A* with B2002 as the increasing parent and *QGN.perg-5A* with the B19 as the increasing parent. Furthermore, the QTL peak of the R5B region was located between 562.0-671.3 Mb (+/− 1 LOD) and harbored *QFS.perg-5B* and *QGLPA.perg-5B.* In this case, the increasing allele was contributed by B2002.

**Table 5.**
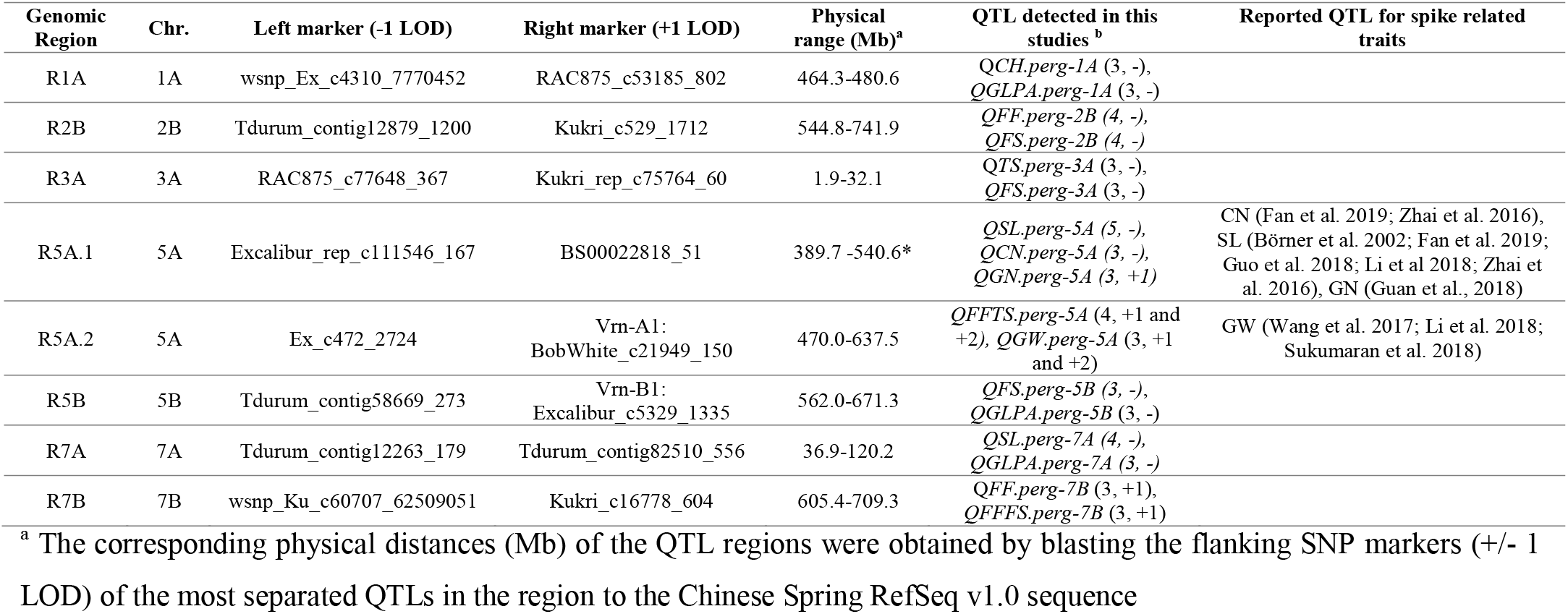
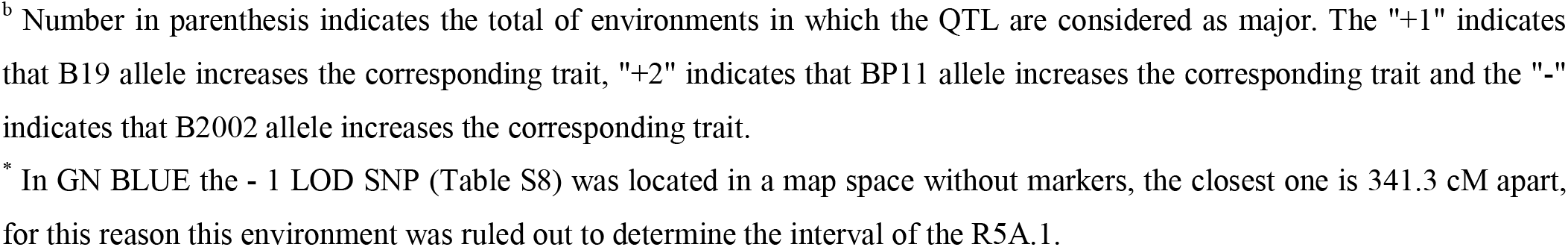
Genomic regions harboring more than one major and stable QTL.

| Genomic Region | Chr. | Left marker (-1 LOD) | Right marker (+1 LOD) | Physical range (Mb)<sup>a</sup> | QTL detected in this studies<sup>b</sup> | Reported QTL for spike related traits |
| --- | --- | --- | --- | --- | --- | --- |
| R1A | 1A | wsnp_Ex_c4310_7770452 | RAC875_c53185_802 | 464.3-480.6 | *QCH.perg-1A* (3, -),<br>*QGLPA.perg-1A* (3, -) |  |
| R2B | 2B | Tdurum_contig12879_1200 | Kukri_c529_1712 | 544.8-741.9 | *QFF.perg-2B* (4, -),<br>*QFS.perg-2B* (4, -) |  |
| R3A | 3A | RAC875_c77648_367 | Kukri_rep_c75764_60 | 1.9-32.1 | *QTS.perg-3A* (3, -),<br>*QFS.perg-3A* (3, -) |  |
| R5A.1 | 5A | Excalibur_rep_c111546_167 | BS00022818_51 | 389.7 -540.6* | *QSL.perg-5A* (5, -),<br>*QCN.perg-5A* (3, -),<br>*QGN.perg-5A* (3, +1) | CN (Fan et al. 2019; Zhai et al. 2016),<br>SL (Börner et al. 2002; Fan et al. 2019;<br>Guo et al. 2018; Li et al 2018; Zhai et<br>al. 2016), GN (Guan et al., 2018) |
| R5A.2 | 5A | Ex_c472_2724 | Vrn-A1:<br>BobWhite_c21949_150 | 470.0-637.5 | *QFFTS.perg-5A* (4, +1 and<br>+2), *QGW.perg-5A* (3, +1<br>and +2) | GW (Wang et al. 2017; Li et al. 2018;<br>Sukumaran et al. 2018) |
| R5B | 5B | Tdurum_contig58669_273 | Vrn-B1:<br>Excalibur_c5329_1335 | 562.0-671.3 | *QFS.perg-5B* (3, -),<br>*QGLPA.perg-5B* (3, -) |  |
| R7A | 7A | Tdurum_contig12263_179 | Tdurum_contig82510_556 | 36.9-120.2 | *QSL.perg-7A* (4, -),<br>*QGLPA.perg-7A* (3, -) |  |
| R7B | 7B | wsnp_Ku_c60707_62509051 | Kukri_c16778_604 | 605.4-709.3 | *QFF.perg-7B* (3, +1),<br>*QFFFS.perg-7B* (3, +1) |  |

<sup>a</sup> The corresponding physical distances (Mb) of the QTL regions were obtained by blasting the flanking SNP markers (+/- 1 LOD) of the most separated QTLs in the region to the Chinese Spring RefSeq v1.0 sequence
<sup>b</sup> Number in parenthesis indicates the total of environments in which the QTL are considered as major. The "+1" indicates that B19 allele increases the corresponding trait, "+2" indicates that BP11 allele increases the corresponding trait and the "-" indicates that B2002 allele increases the corresponding trait.
\* In GN BLUE the - 1 LOD SNP (Table S8) was located in a map space without markers, the closest one is 341.3 cM apart, for this reason this environment was ruled out to determine the interval of the R5A.1.

## 4. Discussion

Most of the breeding progress in wheat yield potential has been achieved by selection of yield *per se* due to the lack of reliable secondary traits and molecular information available to use in MAS (Snape and Moore 2007). The yield potential improvement was, in most cases, consequence of increased GN (Waddington et al. 1986; Perry and D’Antouno 1989; Siddique et al. 1989; Slafer and Andrade 1989, 1993; Acreche et al. 2008; Del Pozo et al. 2014; Lo Valvo et al. 2018), though some recent effects of GW are being reported (Sadras and Lawson 2011; Aisawi et al. 2015; Yao et al. 2019). In the present paper we identified one stable and major QTL for GN in the chromosome 5A (*QGN.perg-5A)* mapping in the same position than the one reported by Guan et al. (2018). However, we recently reported this position as primarily controlling the fertile floret efficiency (FFE, fertile florets per g SDW) when we identified and validated the *QFFE.perg-5A* (Pretini et al. 2020b). Then, as the GN is the result of the FFE and GST (both defining FE) together with the SDW (Fischer 1984, 2011), the *QFFE.perg-5A* can be detected as a QTL associated to GN, highlighting the relevance of FFE and the QTL we validated to define GN. This result exemplifies the importance of dissecting the traits into simpler and more heritable components because it enables a better future search for the actual candidate gen. In relation to GW, we detected two QTL, one on chromosome 5A and other on chromosome 6B (Table 4). The first one has already been reported (Wang et al. 2017; Li et al. 2018; Sukumaran et al. 2018), but the second one, the *QGW.perg-6B* is novel. This QTL is located 157.7 Mb apart from the B genome homeolog (*GW2-B1*) of the *GW2* gene, associated with grain size (Su et al. 2011), suggesting that it would not be a candidate gene to explain the phenotypic variations observed.

The GN is complex a trait itself, being the result of many numerical and physiological spike fertility related traits. In the present study, 25 major and stable QTL for spike fertility related traits were detected (without considering the one for GN and the two of GW already mentioned in the previous paragraph). There were the only two traits for which no stable and major QTL were detected (SDW and GST), which agrees with the low narrow-sense heritability observed (see Table S6) and highlights the high impact of environment on those traits (see Table 3 in Pretini et al. 2020a). For SL, three QTL were detected. The *QSL.perg-2B* is 13.4 Mb apart from a QTL previously described (Cui et al. 2012, Table S1) and the *QSL.perg-5A* is located in the same region as a previously detected QTL (Li et al. 2018; Börner et al. 2002; Fan et al. 2019; Guo et al. 2018; Li et al. 2018; Zhai et al. 2016, Table 1). In contrast, for the *QSL.perg-7A*, no equivalent regions have been detected in previous studies. For TS, three QTL were detected. The *QTS.perg-2D* partially overlaps with a previously detected QTL (Zhou et al. 20170, Table S1). Meanwhile, the *QTS.perg-7A* is in the same region of a previously reported QTL (Cui et al. 2012; Ding et al. 2011; Fan et al. 2019; Jantasuriyarat et al. 2004; Ma et al. 2018; Xu et al. 2014; Zhai et al. 2016, Table 1), and co-localizes with the recently described *WAPO-A1* gene (674.07 Mb) that modifies the total number of spikelet per spike (Kuzay et al. 2019). Finally, the *QTS.perg-3A* detected in our work has not been previously reported. For CN, two major and stable QTL were detected, the *QCN.perg-5A*, which is in the same region as a previous detected QTL (Fan et al. 2019; Zhai et al. 2016), and the *QCN.perg-2A*, which is a novel one. For FF, the two QTL detected are novel. The *QFF.perg-2B* is ~539 Mb apart from the QTL for FF detected by Guo et al. (2017) discarding that it is the same region, while for *QFF.perg-7B*, no equivalent regions have been reported previously. For FS, three QTL were detected. The *QFS.perg-2B* is 540 Mb apart from a QTL detected previously (Deng et al. 2017; Ma et al. 2018, Table 1), ruling out that it was the same region. For the two remaining QTL, *QFS.perg-3A* and *QFS.perg-5B*, regions shared with other works were not detected. For the rest of the traits analyzed in this study (FFTS, FFFS, R, GLPA and CH), no other previous reports are available, to the best of our knowledge (Table S1). Then, we consider that we detected novel QTL for FFTS (*QFFTS.perg-5A* and *QFFTS.perg-5B*), FFFS (*QFFFS.perg-7B*), R (*QR.perg-2A*, *QR.perg-3B* and *QR.perg-6A*), GLPA (*QGLPA.perg-1A*, *QGLPA.perg-3A*, *QGLPA.perg-5B* and *QGLPA.perg-7A*) and CH (*QCH.perg-1A* and *QCH.perg-2B*). No QTL was detected in chromosome 2A for FFTS or FFFS, where the *GNI-A1* gene (Sakuma et al. 2019), known to increase the number of grains through increase of fertile florets per spikelet, has been identified.

As many of the spike fertility traits detected in this study had similar positions, we identified 8 genomic regions that share 17 significant and stable QTL for the different traits (R1A, R2B, R3A, R5A.1, R5A.2, R5B, R7A and R7B). Only in two of these regions (R5A.1 and R5A.2) another QTL for the same trait have previously been described, being the remaining six regions identified for the first time as important hot spots for spike fertility traits (Table 5). Interestingly, the R5A.1 region, which contains significant QTL for SL, CN and GN, is close to the *QFFE.perg-5A* identified and validated for fertile floret efficiency in Pretini et al. (2020b). The allele of B2002 parent increase the SL and CN while the allele from B19 parent increase the GN via *QFFE.perg-5A*. These results agree with the performance of the parental lines described in the present study (Table 3). The region on chromosome 5A (R5A.2), which includes *QFFTS.perg-5A* and *QGW.perg-5A* coincides with the location of the vernalization response gene *Vrn-A1*. While the R5B region, which includes *QFS.perg-5B* and *QGLPA.perg-5B*, coincides with the location of the other vernalization response gene *Vrn-B1*. Li et al. (2019) had recently reported the role of *Vrn1*, together with FUL1 and FUL2 on spikelet and spike development, but the *vrn1* null mutant effect was associated with a high impact on days to heading. The three parental lines of the two DH populations used in the present study are consider spring wheat due to its allelic constitution, *Vrn-A1b*/*vrn-B1*/*vrn-D1* for Baguette 19 and Baguette Premium 11 parents, and *vrn-A1*/*Vrn-B1*/*vrn-D1* for BioINTA 2002, and are mostly insensitive to photoperiod, all parents showed the *Ppd-D1a* allele. This agrees with the anthesis dates of the lines within each population described in Pretini et al. (2020b), which were very close, except for the summer sowing (E5) of B19xB2002 population, where the range was higher. Furthermore, to test the effect of the two functional markers for vernalization (*Vrn-A1* and *Vrn-B1*) we made an ANOVA and detected small differences between the anthesis dates for both populations. For the BP11xB2002 population there was a difference of 3 and 5 day to anthesis depending on the allelic constitution of the *Vrn-A1* and *Vrn-B1* genes, respectively. On the other hand, for B19xB2002 population, depending on the allelic constitution of both genes, there was a difference of 3 day to anthesis. No epistatic interaction was observed between *Vrn-A1* and *Vrn-B1* in the BP11xB2002 population, while a difference of up to 7 days to anthesis was observed depending on the allelic constitution in the B19xB2002 population. Based on that and in the fact that most of the QTL included in the R5A.2 and R5B regions were not expressed in E5 (except for *QGLPA.perg-5B* in B19xB2202) we consider these QTL are not masking an important phenology effect. In contrast, it could be indicating that the *Vrn-A1* and *Vrn-B1* allelic variation in the population might have a pleiotropic effect on the spike traits located in those regions with little impact on phenology in the tested conditions.

The spike fertility related traits are correlated, positively or negatively, depending on the trait (Hay and Kirby 1991). In addition, a negative correlation is usually observed between GN and GW (Slafer and Andrade 1989, Sadras and Lawson 2011, Griffiths et al. 2015). Then, we wonder about the possible pleiotropic effects of each of the 8 regions we detected over the other spike related traits, GW and final yield per spike (YLD), following the Figure 1. For this, we performed an ANOVA for each of the evaluated traits using the highest QTL peak marker as class variables in the model and the environments were included as random class variable. Four regions had a significant effect on GN (R2B, R3A, R5A.1 and R5A.2), six on GW (R1A, R2B, R5A.1, R5A.2, R7A.1, R7A.2, and R7B), but only two in YLD (R5A.1, R5A.2). For the R5A.1 region, where the *QSL.perg-5A, QCN.perg-5A* and *QGN.perg-5A* were located, when the region from B19 is present it results in a shorter spike (−6% SL) without reducing the TS (+2%) or FS (ns), due to a reduction in the distance between spikelets (−5% CN). The FF increases 4% due to higher FFE (+10%), despite a reduction in the SDW (−3%) which is accompanied by a 3% increment of the FFFS. The FF increment together with the higher GST (+8%) results in an increment in GN (+7%), which translate to higher yield (+3%) despite a significant reduction in GW (Figure 1). As we mentioned previously, this region includes the *QFFE.perg-5A* identified and validated for fertile floret efficiency in Pretini et al (2020b), also within the B19xB2002 population and showed similar pleiotropic effects to the R5A.1 region. The other region that resulted in final higher YLD was the R5A.2, which contained the *QGW.perg-5A* and *QFFTS.perg-5A.* When this region from B19 is present the SL is not affected, but the distance between spikelet’s is increased (+3% CN), reducing the TS (−2%). The FFTS and the FFFS increase 3 and 2%, respectively and the FFE is higher (+6%). Nevertheless, the GN is not significantly improved. The YLD improvement of R5A.2 (+5%) when B19 alleles are present is consequence of the increased GW (+6%) (Figure 1). As far as we know, the pleiotropic effect of these region had not been previously reported, expect for Pretini et al. (2020b) for the *QFFE.perg-5A* which is within the R5A.1. We made a similar analysis of pleiotropic effect for each QTL identified, being the *QGW.perg-6B* the only one that has a pleiotropic effect in YLD. When the B2002 alleles are present, the spikes are longer (+2% SL), but the TS and CN is not significantly modified. Nevertheless, higher FS were detected (+2%), which was counterbalanced by a reduction in the FFFS (−2%) resulting in no impact on GN. The YLD increment (+5%) was consequent of the increased GW (+10%). This is an interesting result highlighting the relevance of this QTL for the first time. The complex pleiotropic analysis we preformed, where we analyzed 14 traits and 8 regions, allow as to conclude that the R5A1 and R5A.2 regions together with the *QGW.perg-6B* are of high relevance to be used in MAS to improve a set of traits related with yield potential. All the QTL identified for the spike related traits are the first step to search for their candidate genes, which would allow their better manipulation in the future.

**Fig. 4.**
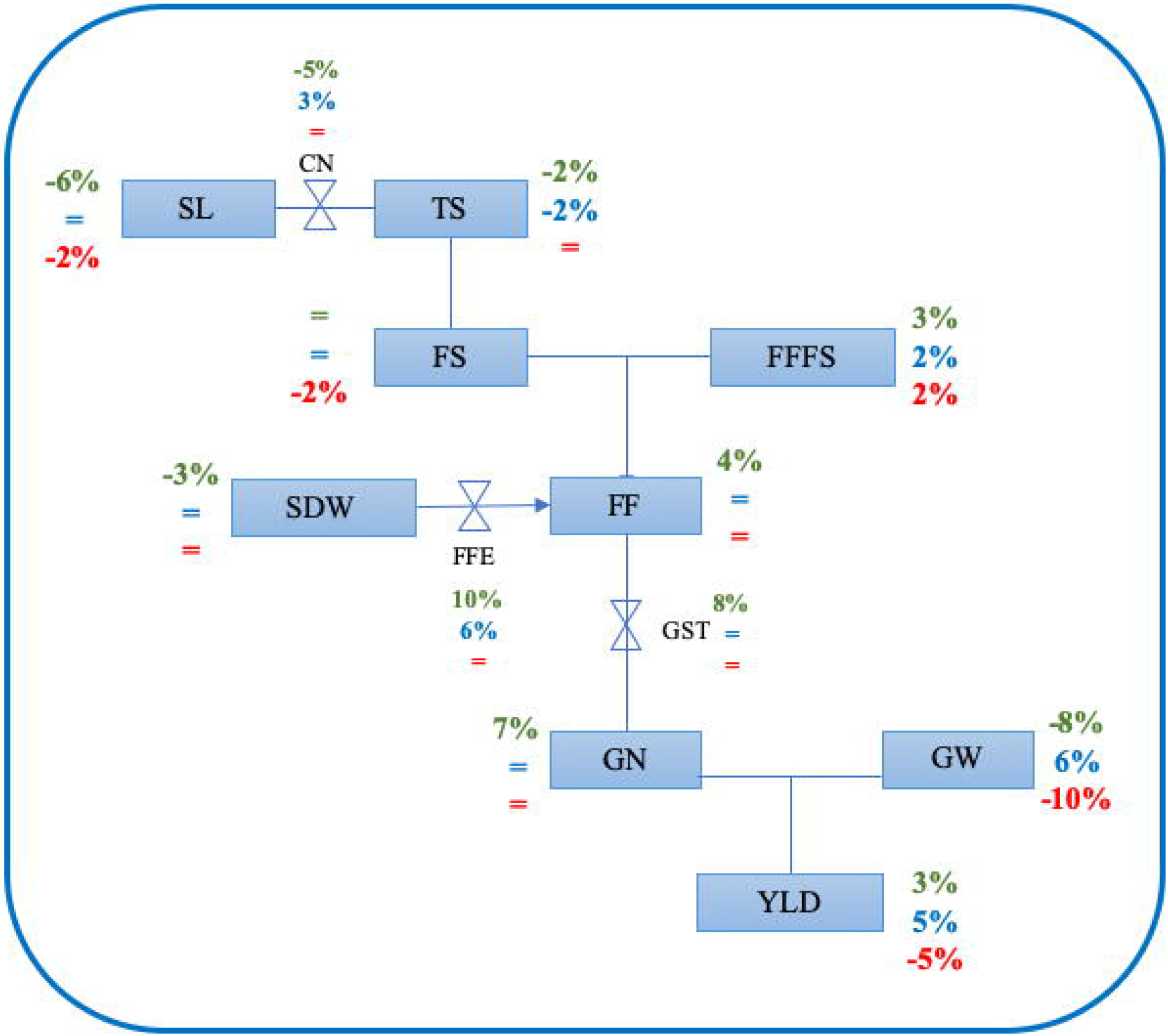
Physiological conceptual framework of analysis of measured variables showing the main and pleiotropic effects of the R5A.1 and R5A.2 regions and *QGW.perg-5A.* The symbols = indicate not significant effect. The green percentage represent R5A.1 while the blue percentage represent R5A.2 and the red percentage represents *QGW.perg-6B.* SL: spike length, TS: total spikelets per spike, CN: compactness of the spike, FF: fertile florets per spike, FS: fertile spikelets per spike, FFFS: fertile florets per fertile spikelet, SDW: spike dry weight at anthesis, FFE: fertile floret efficiency, GN: grain number per spike, GW: grain weight, GST: grain set, YLD: yield.

## Supporting information

ESM_1

## Abbreviations

B19: Baguette 19 B2002 = BioINTA 2002
B2002: BioINTA 2002
BP11: Baguette Premium 11
CH: chaff (no-grain spike dry weight at harvest per spike, mg spike^−1^)
CN: compactness of the spike (mm node^−1^)
DH: double haploid
E1 to E5: testing environments, see Table 2
FF: fertile florets per spike (n° spike^−1^)
FFFS: fertile florets per fertile spikelet (n° spikelet^−1^)
FFTS: fertile florets per total spikelet (n° spikelet^−1^)
FS: fertile spikelets per spike (n° spike^−1^)
GLPA: glume+lemma+palea+awn (mg spike^−1^)
GN: grain number per spike (n° spike^−1^)
GST: Grain set (n° grains floret^−1^)
GW: grain weight (g)
R: rachis (mg spike^−1^)
SDW: spike dry weight at anthesis (mg spike^−1^)
SL: spike length (mm)
TS: total spikelets per spike (n° spike^−1^)

## Acknowledgments

The present work was funded by the Agencia Nacional de Promoción Científica y Tecnológica (PICT 2012-1198, PICT 2014-1283), the Instituto Nacional de Tecnología Agropecuaria (INTA, PNCYO 1127042, 2019-PE-E6-I126-001, 2019-PE-E6-I114-001 Argentina, the Monsanto Beachell-Bourlag Scholarship, the Universidad Nacional del Noroeste de la Provincia de Buenos Aires, (UNNOBA, SIB 2015, SIB 2017, SIB 2019) Argentina and the EU FP7 Funding (ADAPATWHEAT 289842). NP is a research fellow of the Consejo Nacional de Investigaciones Científicas y Técnicas (CONICET) at the Centro de Investigación y Transferencia del Noroeste de Buenos Aires (CITNOBA). We also thank Luis Blanco, Yanel Perez and Octavio Ghio Trebino for the field technical assistance.

## Footnotes

1 https://github.com/juancrescente/lmap

